# Response of adult carrion beetles *Necrodes littoralis* (L.) (Staphylinidae: Silphinae) to selected cadaveric volatile organic compounds: laboratory and field tests

**DOI:** 10.1101/2023.03.02.530784

**Authors:** Anna Mądra-Bielewicz, Joanna Gruszka, Szymon Matuszewski

## Abstract

Carrion insects need to quickly and accurately locate a suitable carcass to maximize their reproductive success. They are attracted by cadaveric volatile organic compounds (VOCs). However, very little is known about VOCs that attract insects at later stages of carrion decomposition. Here, we tested the response of *Necrodes littoralis* (Linnaeus, 1758). (Staphylinidae: Silphinae), a Palearctic beetle that colonizes large carrion late in decomposition, to selected VOCs. First, in the laboratory choice tests we demonstrated that the beetles reveal no preference for meat with larval blow flies over meat alone. This finding indicates that both, the fly larvae and the feeding matrix they form on meat are not the source of specific attractants for the adult beetles of *Necrodes* Leach, 1815. Therefore, we focused on VOCs that are related to carrion putrefaction. We tested the response of the beetles to benzyl butyrate, butan-1-ol, butyric acid, cadaverine, dimethyl disulfide (DMDS), dimethyl trisulfide, indole, phenol, putrescine and skatole in laboratory choice assays and field trapping tests. None of the compounds elicited the positive and significant response of the beetles under laboratory or field conditions, indicating that these VOCs are probably not the attractants of *N. littoralis*. Moreover, in the field tests we found a significant attraction of *Lucilia sericata* (Meigen, 1826) (Diptera: Calliphoridae) to traps with DMDS. DMDS revealed also a positive (however insignificant) response of *Saprinus* spp. (Coleoptera: Histeridae) and *Sarcophaga* spp. (Diptera: Sarcophagidae). *Sarcophaga* flies were also attracted to traps with butyric acid. These findings expand the knowledge on chemoecology of carrion insects, highlighting the need to further search for VOCs that attract late-colonizers of carrion.

---

Carrion is used by various insects that depend on it in different ways (1). Some are obligate necrophages that can only reproduce on carrion. The location of a suitable carcass is critical for them. In most cases carcasses are distributed in a patchy way (2, 3). Their occurrence is spatially unpredictable, and for this reason carrion insects must rely on olfactory systems, which are effective over long distances. Moreover, carrion is frequently bulky, highly nutritious and ephemeral, causing intense competition among its users (4, 5). Carrion insects need to locate it quickly to maximize the chance for reproductive success. Due to these constraints, obligate necrophages had to develop olfactory systems linked with certain cadaveric volatile organic compounds (VOCs) that enabled them to quickly and accurately locate carrion.

VOCs that attract carrion insects were the focus of many studies in carrion ecology and forensic entomology (6). Sulfur compounds that are released early after death, e.g. dimethyl disulfide (DMDS) or dimethyl trisulfide (DMTS) were demonstrated to attract several species of blow flies (Calliphoridae) and silphid beetles (Staphylinidae: Silphinae) (7-11). These compounds are also produced by carrion mimicking plants to lure carrion insects that are ultimately used by the plants for pollination (12). Since sulfur compounds attract mainly insects that colonize carrion early in decomposition, these VOCs can be classified as universal fresh carrion cues. However, there is a regular succession of different insect groups during carrion decomposition (13) that follows changes in the carrion quality (14), and insects that colonize carcasses later in decomposition are probably attracted by different VOCs than early colonizers. To get the full chemical picture of the necrobiome, these compounds need to be identified and their links with particular insect species need to be studied.

*Necrodes littoralis* (Linnaeus, 1758) (Staphylinidae: Silphinae) is a carrion beetle that colonizes large cadavers in natural (open or forest) habitats of the Palearctic region (15-18). Adult beetles of *Necrodes* Leach, 1815 abundantly appear on carrion near its peak bloating, females oviposit into the soil nearby and the necrophagous larvae form large aggregations late in decomposition (16, 19-21). Upon arrival on carrion adult beetles clear it of the blow fly larvae (4, 19). While feeding in aggregations, *Necrodes* larvae form the feeding matrix, in which carrion substrate is exodigested and heat is produced to benefit the larvae (22). *Necrodes littoralis* also colonizes human cadavers and for this reason the species is useful for the post-mortem interval estimation with the methods of forensic entomology (15, 23-26).

Since no previous study has addressed the question about VOCs that attract *N. littoralis* to carrion, we decided to perform laboratory choice tests and field trapping tests to identify the attractants of these beetles. Adult *Necrodes* beetles were regularly observed to feed voraciously on blow fly maggots (4, 19), so we wanted to first test the hypothesis that these beetles are attracted to VOCs released by larval blow flies or their feeding environment. For this purpose we investigated, whether the beetles reveal a preference for meat with feeding blow fly larvae over meat without the insects. Then, we tested the hypothesis that the beetles are attracted by some VOCs released during carrion putrefaction. We studied response of the beetles under laboratory and field conditions to several VOCs produced during carrion decomposition. Although our field trials were designed to identify attractants of *N. littoralis*, the traps used for this purpose were also efficient in catching other insects. Hence, the study provided also data on the response of other necrophilous insects to the tested VOCs.

## Results and discussion

First, in the laboratory choice tests we found no preference of adult *N. littoralis* for meat with larval blow flies over meat alone (Wilcoxon test: *Z*=0.3; *p*=0.76; Fig. 1 and supplementary Fig. I). *Necrodes* beetles were regularly found to intensively feed on blow fly maggots (4, 15, 19). This is usually considered as predatory behavior (27). However, a recent analysis indicated that this behavior is more related to the competition with blow flies over carrion resources and less to the nutrition of the beetles. Laboratory behavioral assays demonstrated that the beetles highly preferentially killed blow fly larvae that were just before or at the beginning of their peak feeding phase (i.e. late second instar and early third instar larvae), while the more nutritious post-feeding larvae were killed four times less frequently (4). Moreover, the reproductive success of the beetles was the highest when there were no or very few blow fly larvae on a cadaver (4). Therefore, blow fly larvae are not the food target for adult *Necrodes* beetles, although when they are present on carrion the beetles highly effectively eliminate them. Since we found no statistically significant preference of the beetles for the meat with blow flies compared to the meat alone and current study design would demonstrate a potential effect assuming a size of 0.6 or more (see supplementary 1 for more details), the fly larvae or the feeding matrix they form on meat are highly probably not the source of specific attractants for adult *Necrodes* beetles. In effect, we focused on VOCs that are related to carrion putrefaction.

**Fig. 1.**
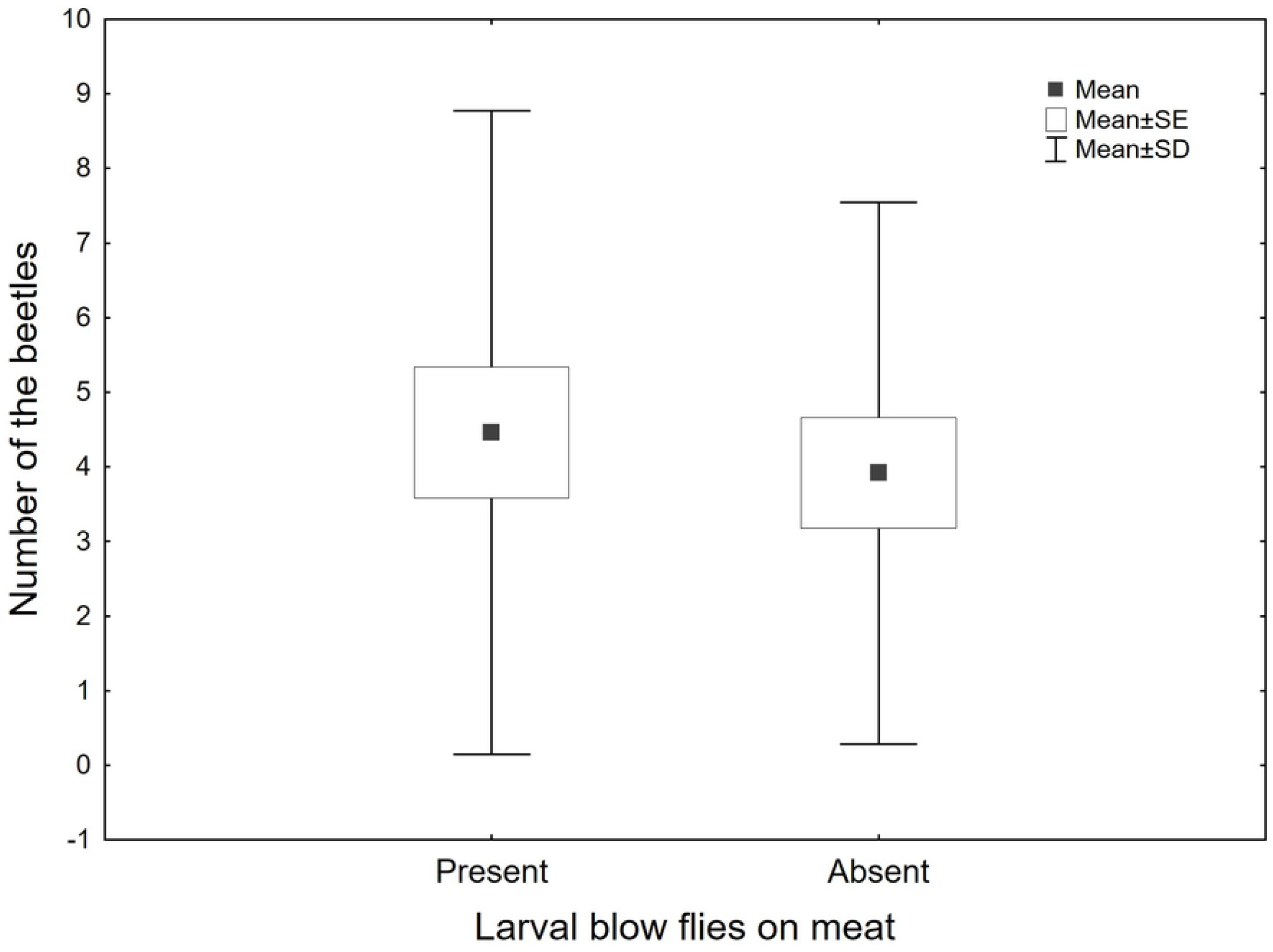
Response of adult *Necrodes littoralis* in laboratory choice tests to the meat with and without larval blow flies. Upon termination of a trial we counted beetles present on meat and in a nearby soil in the experimental and control containers. SE: standard error of the mean; SD: standard deviation.

In the laboratory choice tests with single VOCs no compound elicited a significant positive response of the beetles, phenol was close to significance (*p*=0.09; Table 1; supplementary Fig. II-VII). Dimethyl disulfide (DMDS) and cadaverine revealed a close to significant negative response of the beetles, i.e. more of them were collected in the control area, suggesting that they avoid DMDS and cadaverine (*p*=0.14 and 0.21, respectively; Table 1, supplementary Fig. III and VI). A combination of compounds that, in a single VOC tests, elicited positive responses revealed a statistically significant negative effect (Wilcoxon test: *Z*=2.3; *p*=0.02; Fig. 2). In the field tests we trapped no single specimen of *N. littoralis* in any of the setups, which demonstrates that under field conditions concentrations and combinations of VOCs that were used in these studies do not attract adult *N. littoralis* (Tables 2-3, supplementary Tables I-III and Fig. VIII). As a whole, the current study indicates that benzyl butyrate, butan-1-ol, butyric acid, cadaverine, DMDS, dimethyl trisulfide (DMTS), indole, phenol, putrescine and skatole are probably not the attractants for *N. littoralis*. Although the number of replicates in the laboratory and field tests with VOCs may seem small, it should be evaluated based on the size of the potential effect (see supplementary Fig. VII-VIII). In general, silphid beetles are highly efficient in locating carrion, in spite of its patchy and spatially unpredictable nature (20). In order to quickly and accurately locate the proper carrion, they need to be highly sensitive to the VOCs that are characteristic for such resources. As a result, we predicted that we would see large effects if the relevant VOCs were studied and we decided to study more compounds but with fewer replicates. Although in the field tests the relatively small quantities of chemicals were used, we maximized the chance of a positive beetle response to the VOCs by using the beetle release setup. Moreover, these quantities were sufficient to lure other necrophilous insects, particularly in the beetle release setup. The significant repellent effect of the combination of compounds that elicited positive responses in a single VOC tests is difficult to interpret. The beetle behavior in this case could be an effect of the relatively high amount of the mixture used or the synergistic repellent effect of some VOCs’ combinations.

**Table 1.** Significance (Wilcoxon test) of adult *Necrodes littoralis* response in the laboratory choice tests to selected volatile organic compounds (VOCs).

| VOC | valid N | Z | P |
| --- | --- | --- | --- |
| Benzyl butyrate | 9 | 0.059 | 0.953 |
| Butanoic acid (butyric acid) | 10 | 0.311 | 0.756 |
| Butan-1-ol | 10 | 0.102 | 0.919 |
| Butane-1,4-diamine (putrescine) | 9 | 0.829 | 0.407 |
| Dimethyl disulfide (DMDS) | 10 | 1.478 | 0.139 |
| Dimethyl trisulfide (DMTS) | 10 | 0.408 | 0.683 |
| Indole | 10 | 0.204 | 0.838 |
| 3-methylindole (skatole) | 9 | 0.829 | 0.407 |
| Pentane-1,5-diamine (cadaverine) | 9 | 1.244 | 0.214 |
| Phenol | 8 | 1.680 | 0.093 |

**Fig. 2.**
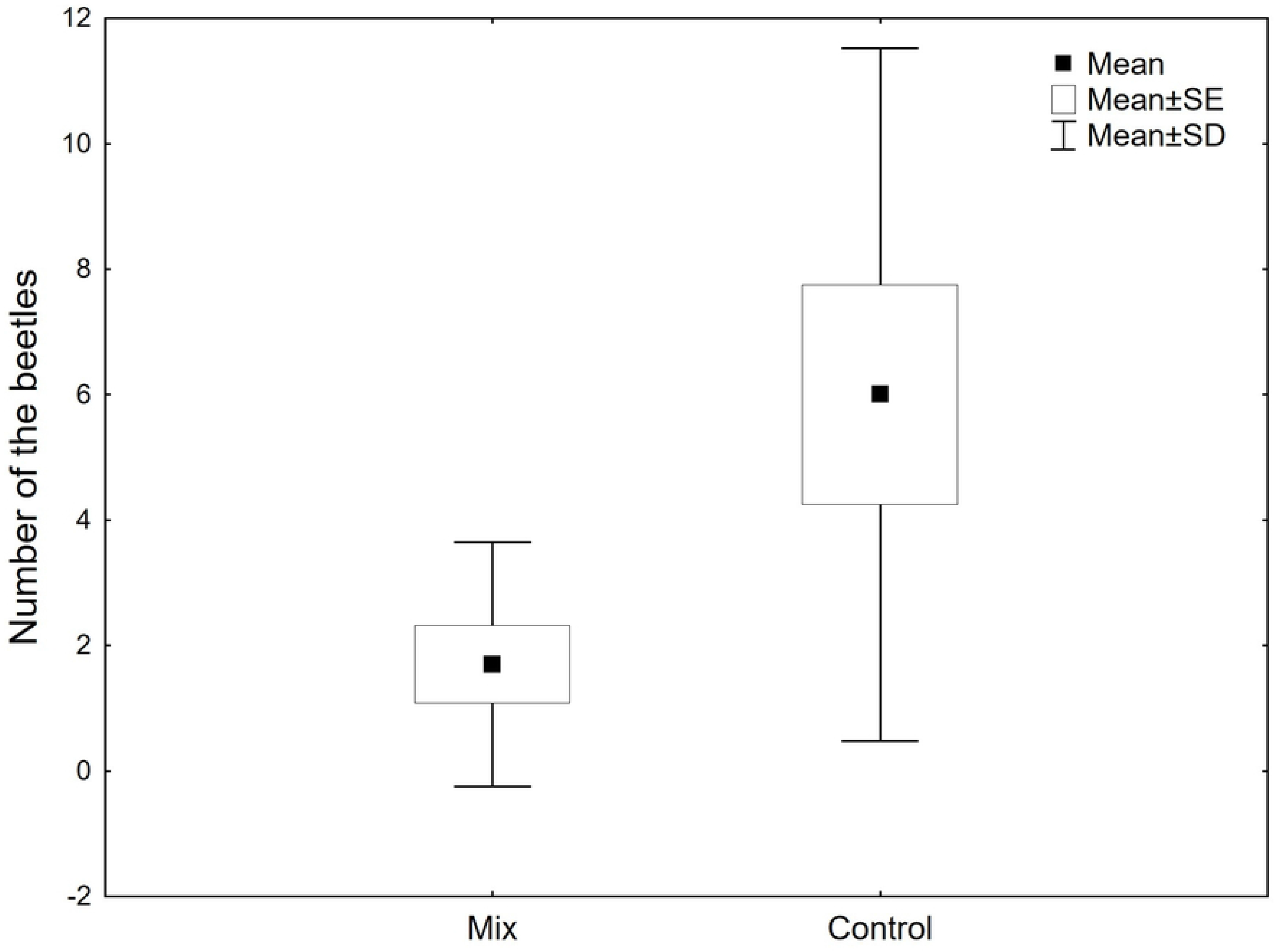
Response of adult *Necrodes littoralis* to the combination of compounds that elicited a positive response of the beetles in the single-compound choice tests (i.e. benzyl butyrate, butyric acid, butan-1-ol, putrescine, dimethyl trisulfide and phenol). Upon termination of a trial we counted beetles present on and in soil in the experimental and control containers. SE: standard error of the mean; SD: standard deviation.

**Table 2.**
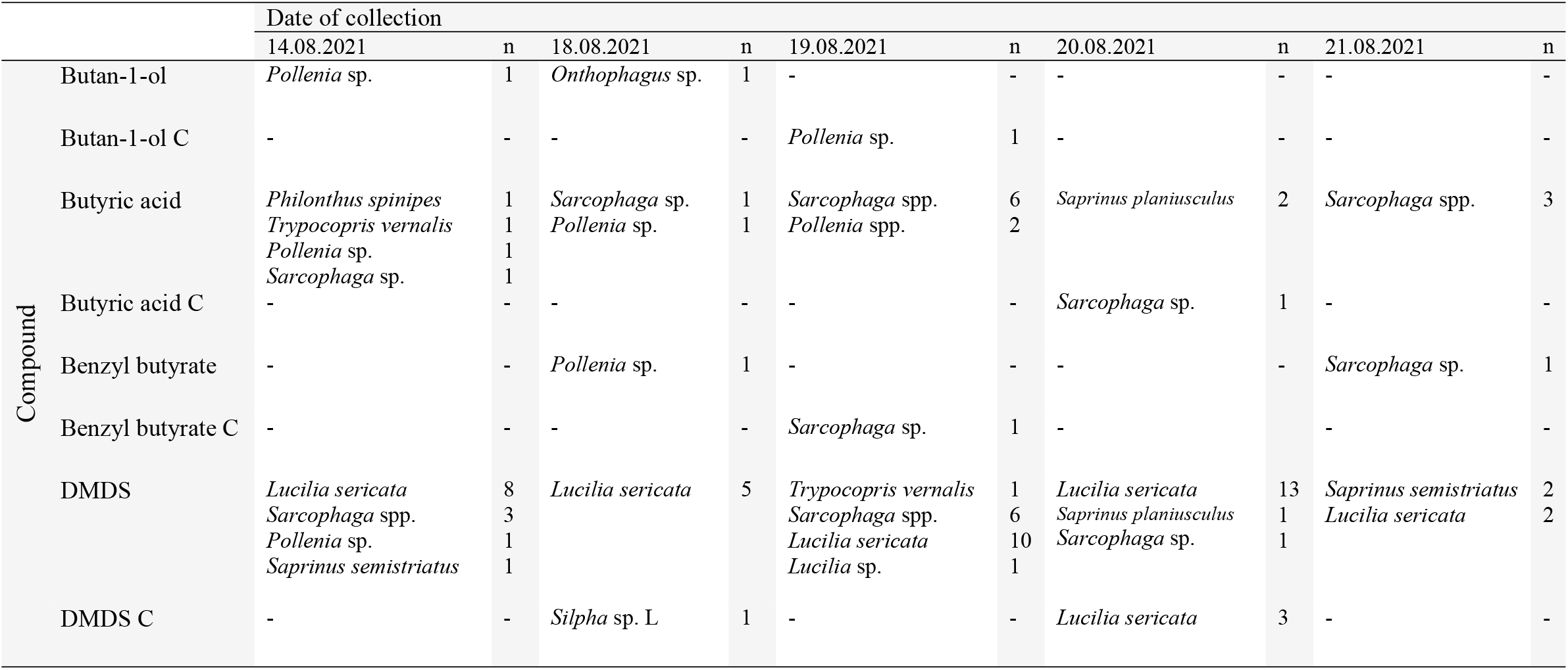
Necrophilous insects trapped during field trials with the release of the beetles using butan-1-ol, butyric acid, benzyl butyrate and dimethyl disulfide (DMDS) as attractants (C – control trap, L – larva).

**Table 3.**
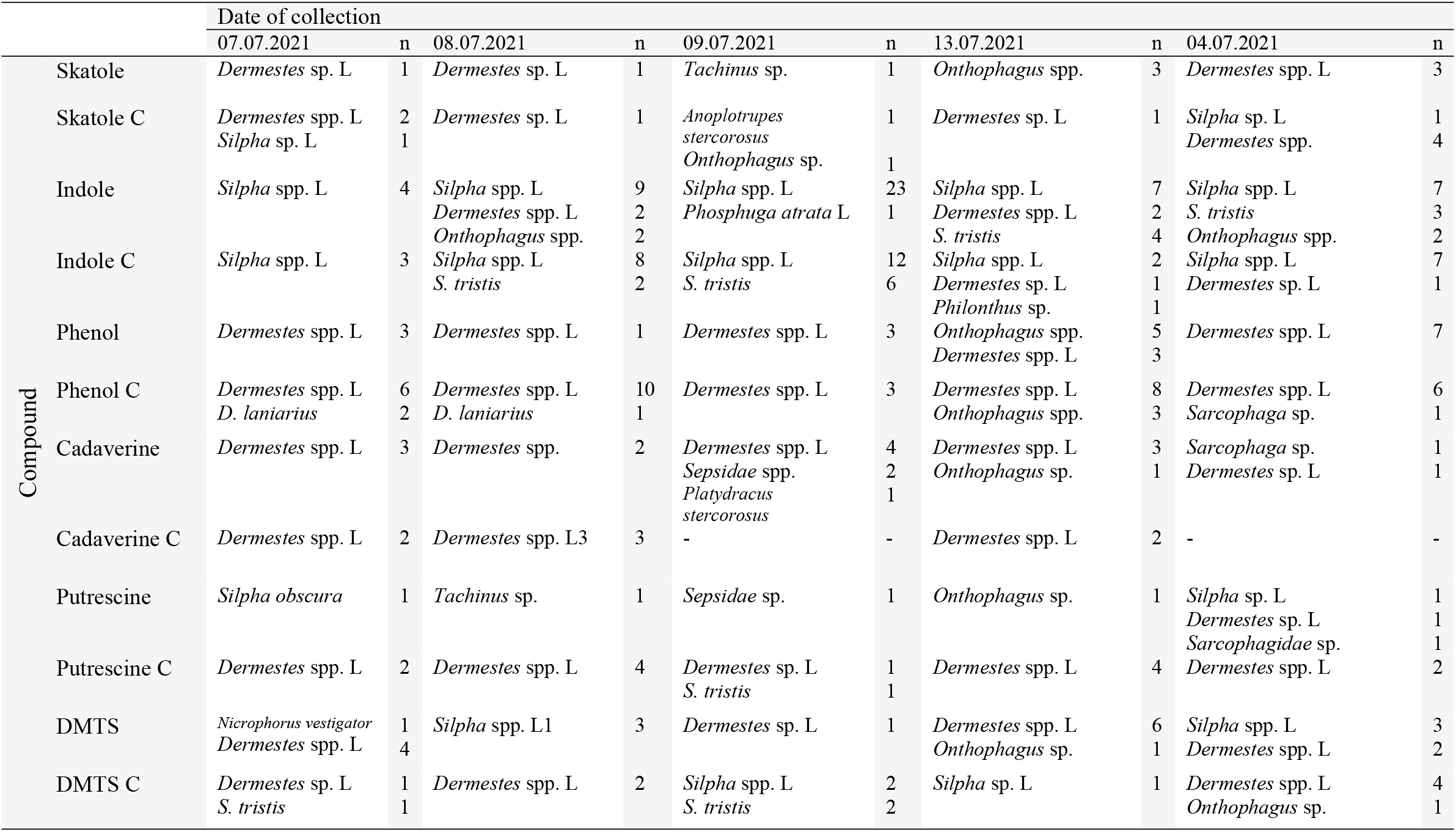
Necrophilous insects trapped during field trials with the release of the beetles using skatole, indole, phenol, putrescine, cadaverine, and dimethyl trisulfide (DMTS) as attractants (C – control trap, L – larva).

The question about attractants of *Necrodes* beetles remains open. We believe that *N. littoralis* is attracted to carrion by some VOC, which simply has not been used in this study. However, we accept the small chance that too small concentrations or wrong combinations of the VOCs used in this study were the reason for the negative results reported in this article. Not surprisingly, we found no response of the beetles to the organosulphur compounds (i.e. DMDS and DMTS). These chemicals are byproducts of microbial decomposition of amino acids that contain sulphur (i.e. cysteine and methionine) and are released from cadavers shortly after death (28-32). Sulfur compounds were found to attract various species of carrion insects including *Lucilia* spp., *Calliphora* spp., *Protophormia terraenovae* (Robineau-Desvoidy, 1830) (Diptera: Calliphoridae), *Thanatophilus sinuatus* (Fabricius, 1775), *Nicrophorus* spp. (Coleoptera: Staphylinidae: Silphinae) or *Nasonia vitripennis* (Walker, 1836) (Hymenoptera: Pteromalidae) (7-12, 33, 34). Sulphur compounds were actually the most frequently reported attractants of carrion insects (6). However, most of the insects that positively responded to these compounds colonize carrion early after death (35-37). In contrast, *Necrodes* beetles are late colonizers (15, 38). Therefore, their response to VOCs emitted early during decay would be a surprise.

While choosing compounds to be used in the experiments, we expected that the attractant of *N. littoralis* was a carrion-originating VOC, which is released later during decomposition as a byproduct of putrefaction (e.g. cadaver related amines). For the tests we also chose the compounds that were found to attract other insects colonizing carrion late in decomposition, e.g. benzyl butyrate that elicited a positive response of *Dermestes maculatus* De Geer, 1774 (Coleoptera: Dermestidae) (39). Unfortunately, none of the tested VOCs (Tables 1-3) gave a positive response of *N. littoralis* in laboratory or field trials. The time when adult *N. littoralis* start to visit pig carcasses is positively correlated with the onset of cadaver bloating (40). Bloating is the distension of carrion that results from the accumulation of gasses produced by bacteria during putrefaction (41). Some component of these gasses probably attracts *Necrodes* beetles; nevertheless, we know very little about VOCs eliciting bloating. Therefore, future studies should characterize gasses responsible for carrion bloating to reduce the set of compounds for choice tests of *Necrodes* beetles. Another possibility is related to quorum sensing signals used by bacteria to communicate with each other (42). It has been suggested that carrion insects may use these signals to locate proper carrion resources (43, 44). Here, we tested indole – an environmental cue used by different organisms for various purposes (45). The beetles did not respond to this compound, however other compounds connected to quorum sensing of carrion bacteria should be the target for future research.

We found a significant attraction of *Lucilia sericata* (Meigen, 1826) (Diptera: Calliphoridae) to the traps with 8 ml of DMDS (Wilcoxon test: *Z*=2.02; *p*=0.043; Fig. 3; Table 2, supplementary Fig. VIII), close to the significant attraction of *Saprinus* spp. (Coleoptera: Histeridae; Wilcoxon test: *Z*=1.60; *p*=0.109; supplementary Fig. VIII-IX; Table 2) and *Sarcophaga* spp. (Diptera: Sarcophagidae; Wilcoxon test: *Z*=1.60; *p*=0.109; supplementary Fig. VIII-IX; Table 2) to these traps and *Sarcophaga* spp. to traps with 8 ml of butyric acid (Wilcoxon test: *Z*=1.48; *p*=0.138; supplementary Fig. VIII and X; Table 2). Individual specimens of other necrophilous taxa were collected in traps with DMDS, DMTS and butyric acid (Tables 2-3, and supplementary Tables I-III).

**Fig. 3.**
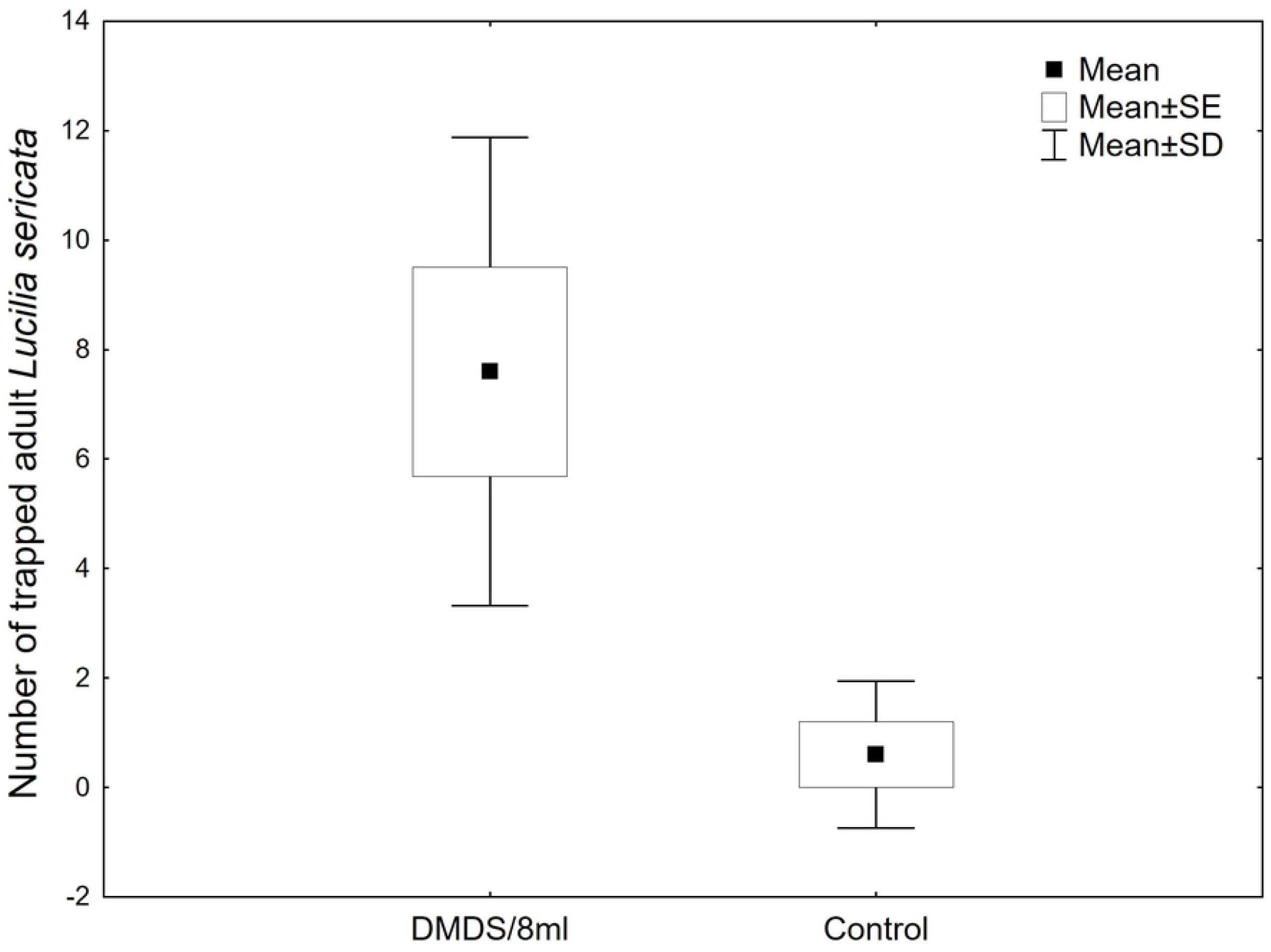
Attraction of adult *Lucilia sericata* to the field traps with 8 ml of dimethyl disulfide (DMDS). SE: standard error of the mean; SD: standard deviation.

These findings confirm the importance of sulphur compounds as attractants of carrion insects. It has previously been demonstrated that *L. sericata* responds to DMDS (8, 46), although there are also data indicating that DMTS is a primary attractant for this species (47, 48). Current experiments revealed the clear response of *L. sericata* to DMDS under field conditions and no response to DMTS. As these compounds were not tested in the field at the same time and in the case of DMTS rather small concentrations were used, our findings simply confirmed importance of DMDS for *L. sericata* attraction to carrion, highlighting at the same time variation in the blow fly response to experimentally released VOCs under field conditions. The negative result for DMTS traps should be therefore interpreted with caution. Interestingly, early field studies indicated that *L. sericata* responds also to hydrogen sulphide (46). Taken together, these data suggest that all carrion-originating sulphur compounds can attract *L. sericata* to carrion, and the fly response is likely highly dependent on the amount of a compound rather than on its specific kind. Current field data demonstrate for the first time that adult *Sarcophaga* and *Saprinus* spp. also respond to DMDS. These insects colonize cadavers early in the succession (35, 49-51). Therefore, their response to DMDS confirms the importance of sulfur compounds in attracting carrion insects, but again early-colonizers. *Saprinus* beetles are predators of fly larvae, particularly blow flies (50). Their response to DMDS, a compound released both by the carrion itself and by the third instar larvae and pupae of blow flies (52), may be an example of attraction by the prey’s environment or the prey itself.

In conclusion, this study demonstrated that *N. littoralis* beetles reveal no preference for meat with larval blow flies over meat alone. Moreover, we found no positive and significant response of the beetles under laboratory or field conditions to benzyl butyrate, butan-1-ol, butyric acid, cadaverine, DMDS, DMTS, indole, phenol, putrescine and skatole, indicating that these VOCs are probably not the attractants of *N. littoralis*. There are several limitations of this study, the most important are the relatively small amounts of VOCs and low number of replicates used in the field tests as well as the small number of compound combinations included in the study. Therefore, current negative findings regarding particular VOCs should be interpreted cautiously.

## Materials and methods

### Insects

Beetles used in the experiments came from our main laboratory colony (selected at random). The colony was established in June 2017 and replenished in June 2018 using beetles collected in alder forest near Biedrusko (Western Poland, CE, 52°31’N, 16°54’E). Males and females were kept separately in 7.5 l terrariums (20-40 beetles per terrarium, at least four terrariums at the same time), on soil, with pork meat *ad libitum* and cotton wool soaked with water. Beetles used in the laboratory tests were about one month old, after completing a trial they were returned to the main colony. In the subsequent trials the same pool of the beetles was used. Beetles released in the field tests were about two weeks old.

### Laboratory choice tests

In the initial trials behavior of the beetles was tested in the two-arm olfactometer (a central rectangular arena with two arms of about 50 cm made of PVC [polyvinyl chloride] and with fans at the endpoints of the arms, supplementary Fig. XI). The device was positioned on a laboratory table, with no extra barrier to the background volatiles and no filtration of the air used by the fans. We analyzed the response of the beetles to meat samples of various degrees of decay (1, 2, 3 and 4 days old pork samples against each other and against sterile cotton swabs). Insects were tested in various light conditions (with different intensities and colors of artificial light) and also in a separate room (complete dark and no contaminant VOCs from the background). In all of these conditions behavior of the beetles in the olfactometer was erratic and difficult to interpret. Some insects swiftly chose one arm and ran through it without any clear searching behavior or visible interest in the olfactory stimuli, whereas the others stayed immobile in the central arena for a long time. Although several modifications of the setup were tested, we failed to induce the typical searching behavior of the beetles. Therefore, we decided not to use the olfactometer setup in further trials and not to present the data from olfactometer assays in this article, and instead to test behavior of the beetles under less stressful conditions (as described below).

Choice trials were conducted in Exo Terra 46 × 30 × 17 cm plastic terraria, filled to 1/2 with a humid substrate for potted plants. Two smaller containers (15 × 10 × 6.5 cm) were put at the opposite walls of the terrarium and were fully covered with the substrate (Fig. 4). These test and control containers were used to facilitate the sampling of the beetles upon termination of a trial. The setup was used to test the response of the beetles to the meat with larval blow flies and to the VOCs.

**Fig. 4.**
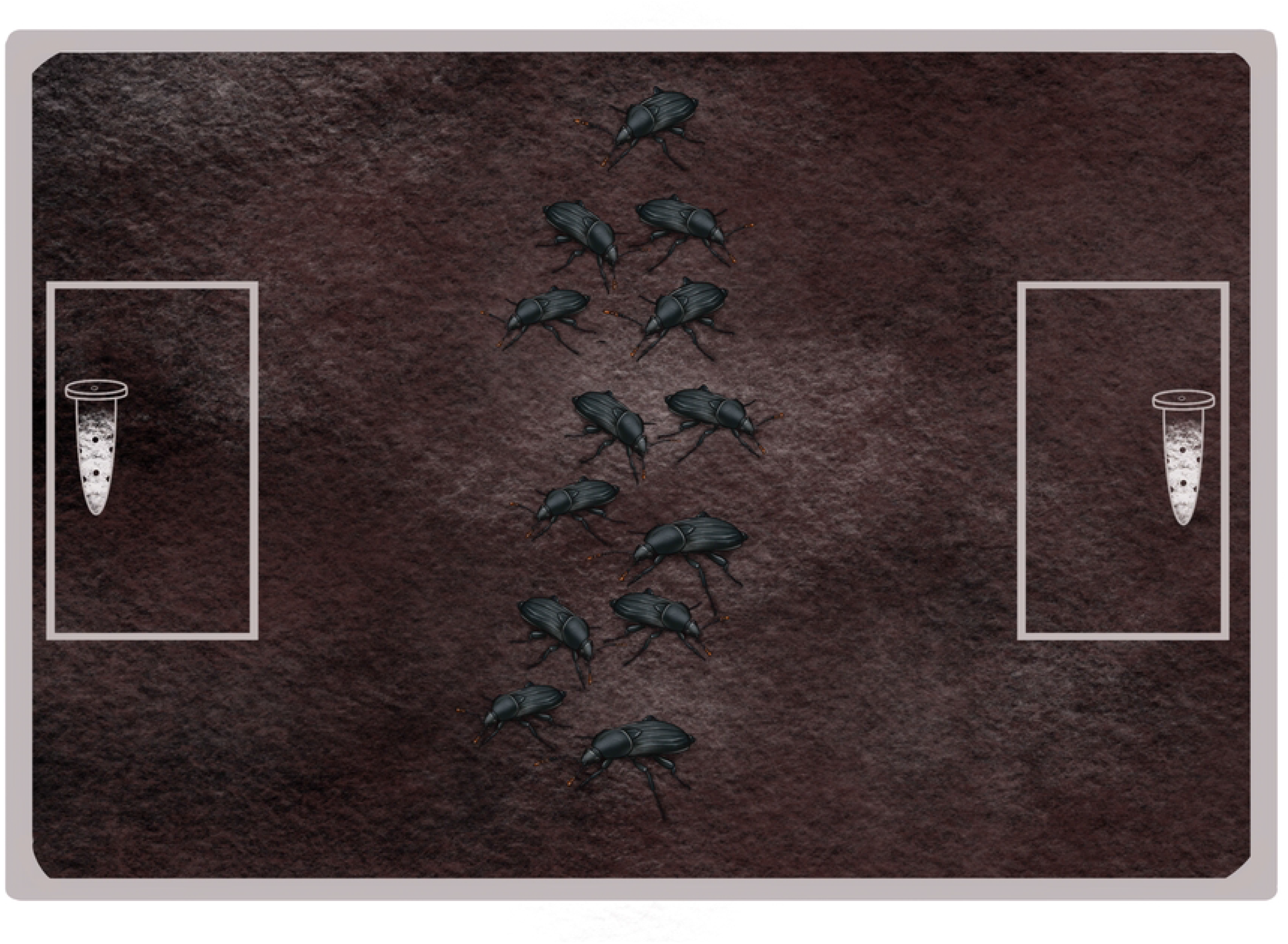
The setup used in the laboratory choice tests of *Necrodes littoralis* with VOCs. Experimental and control Eppendorf tubes were positioned on soil at the opposite ends of a large terrarium. Upon termination of a trial smaller experimental and control containers were taken out and beetles present inside were counted.

In the case of the experiments with larval blow flies, we used larvae of *Calliphora vomitoria* (Linnaeus, 1758) (Calliphoridae). They were reared on pork in 500 ml plastic containers with a perforated lid, on a thin layer of soil, in a temperature chamber (ST 1/1 +, POL EKO, Poland) at 20 °C and photoperiod 12:12 (L/D). We used late second instar or early third instar larvae only, since they are preferentially killed by adult *Necrodes* beetles (4). Fifty larvae were put on a piece of pork meat that was partially wrapped in an aluminum foil. The control piece of meat was also partially wrapped in an aluminums foil. Both pieces were put for an hour into the temperature chamber and then they were transferred to the opposite sides of the main container in the middle of the test or control containers. Test and control sides of the terrarium were swapped after each trial. The whole setup was covered with aluminum foil to minimize the effects of stressors on the beetles. Terraria were kept under the fume hood at room temperature and humidity (20-23°C, 50-60%). Forty adult *N. littoralis* (20♀ and 20♂) were put inside a terrarium at about 1 p.m., next day at 8 a.m. smaller containers were taken out of the terrariums and we counted beetles present inside the soil substrate, on its surface or on meat in test or control containers and outside of them. The experiment was replicated 25 times.

In the case of the experiments with VOCs test and control samples were provided to the beetles in Eppendorf tubes (1.5 ml) with 7 needle holes and cotton wool inside (Fig. 4). Test samples contained 0.5 ml of the given chemical; control samples contained only cotton wool. Eppendorf tubes were positioned on the soil substrate in the centers of the smaller containers. Then adult beetles were added following the protocol used in the choice tests with larval blow flies. Ten compounds were tested (supplementary Table IV). A single compound was tested at the same time, after completing trials with the given compound, the substrate inside terraria was replaced. The trials with each compound were replicated 10 times. After trials with single compounds, we studied the response of the beetles to the combination of chemicals that elicited a positive response of the beetles in the single-compound trials (i.e. benzyl butyrate, butyric acid, butan-1-ol, putrescine, DMTS and phenol, 0.25 ml of each).

The significance of differences (at α=0.05) in the number of beetles present in control and test containers used in the choice tests was evaluated using the Wilcoxon signed-rank test (Statistica 14, TIBCO Software Inc.). Post-hoc power analysis was performed using G*Power 3.1 (53).

### Field tests

We used setups without and with the release of the beetles in the field. In both cases pitfall traps (a plastic pot with water and a roof, Fig. 5) were buried along a straight line every 30 m. Experiments were performed on Campus Morasko of Adam Mickiewicz University in Poznań (Western Poland, 52°469’N, 16°918’E) in a meadow habitat, sparsely covered with shrubs and single trees.

**Fig. 5.**
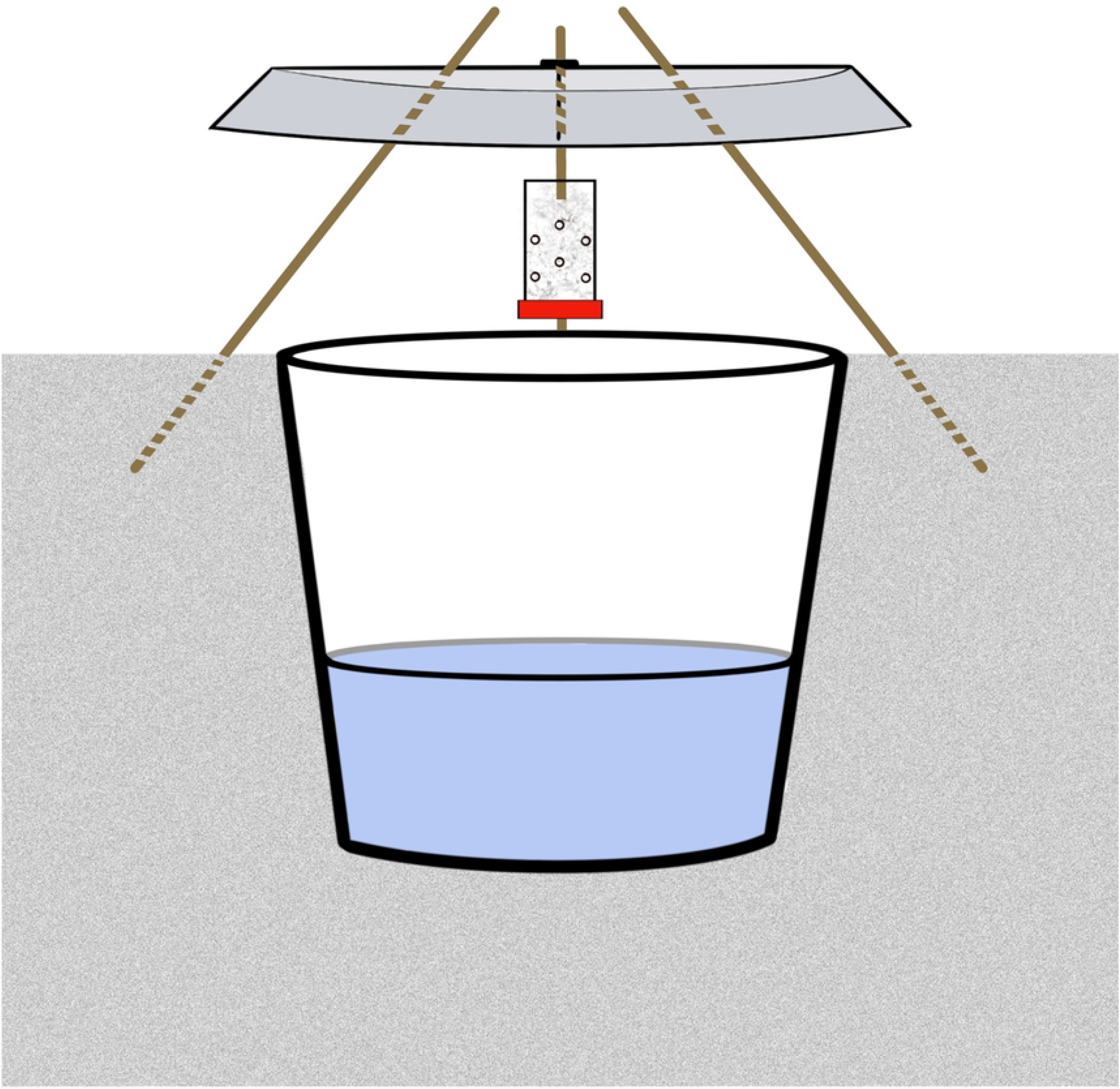
Construction of the pitfall trap used in the field tests. We used a plastic pot (volume: 2.1 l., top ø: 17cm) and a roof (plastic plate) attached to the ground with wooden kitchen sticks. A 60 ml plastic container with 10 needle holes, a cotton wool and a tested compound was suspended under the roof. The traps were 1/3 filled with water (with the addition of soap to diminish surface tension).

In the case of experiments without the release of the beetles, we used two setups. In the SETUP1 every experimental trap had a corresponding control trap (about 10 m away), in the SETUP2 we used two control traps at the ends of the line formed by experimental traps. With the SETUP1 we used Eppendorf tubes (1.5 ml) with 7 needle holes, cotton wool inside and 1 ml of a compound. The tubes were suspended under the roof in the center of the trap. This setup was used to test DMDS, benzyl butyrate, butyric acid and butan-1-ol. Ten trials were made from May 19 until June 1 2020. In the SETUP2 we modified the carrier of attractants to enable using more chemicals on a single carrier. Instead of Eppendorf tubes we used cotton tampons with 2, 4, 6 and 8 ml of chemicals to study one, two, three or four compounds, respectively. We tested DMDS, benzyl butyrate, butyric acid and butan-1-ol from June 4 until June 18 2020 (10 trials), combinations of DMDS with benzyl butyrate, butyric acid or butan-1-ol from June 4 until July 24 2020 (20 trials), a combination of DMDS with benzyl butyrate and butan-1-ol and a combination of all four compounds from July 8 until July 24 2020 (10 trials). In both setups, the traps were emptied and attractant carriers were replaced daily at noon.

In the case of experiments with the release of the beetles, every experimental trap had a corresponding control trap (about 10 m away, Fig. 6). To further increase the amount of chemicals on a single carrier and lengthen their release time, we used 60 ml plastic containers with 10 needle holes, cotton wool and 8 ml of a compound. The containers were suspended under the roof in the center of the trap. This setup was used to test DMDS, benzyl butyrate, butyric acid, butan-1-ol (5 trials, from August 14 until August 21, 2020), putrescine, cadaverine, indole, skatole, phenol and DMTS (5 trials, from July 7 until July 14, 2021). At each pair of experimental and control traps we released 10 adult beetles (5♀,5♂, the beetles were not marked in any way) at a release point approximately 60 m away from the traps (Fig. 6). Carriers with attractants were put in the field at 2 p.m., beetles were released at 4 p.m. the same day and the next day traps were emptied at 10 a.m.

**Fig. 6.**
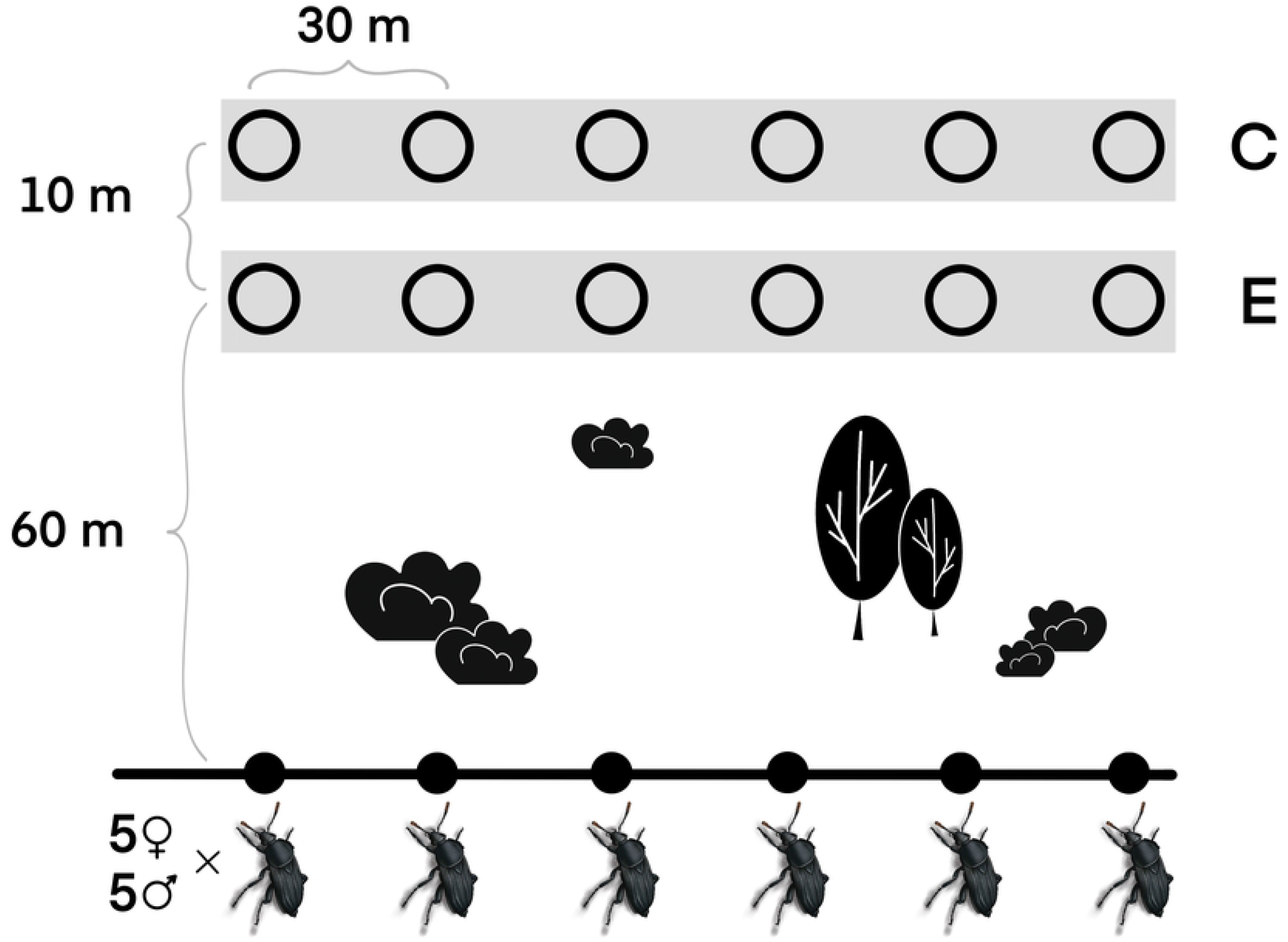
The setup used in the field tests with the release of the beetles. C: control traps, E: experimental traps

We identified to the species level most of the necrophilous insects. The significance of differences (at α=0.05) in the number of trapped specimens of a given species between experimental and control traps was evaluated using the Wilcoxon signed-rank test (Statistica 14, TIBCO Software Inc.). Post-hoc power analysis was performed using G*Power 3.1 (53).

## Ethical approval

Laboratory and field tests used an insect species *Necrodes littoralis* (Linnaeus, 1758) (Staphylinidae: Silphinae). The species is not under protection. No permission or approval from Ethics Commission was needed.

## Data availability

The datasets supporting this article are available from the corresponding author on a reasonable request.

## Acknowledgments

The study was funded by the National Science Centre of Poland (grant no. 2016/21/B/NZ8/00788). We thank anonymous reviewers for their comments that helped us to improve the manuscript.

## Author contributions

S.M. developed the concept for the study and the article, analyzed the data, prepared the graphs and wrote the manuscript. A.M.B. identified insects sampled in the field tests, drew the figures and prepared the tables with results of the field trials. All authors performed experiments, prepared raw data for the analyses, discussed the results and reviewed the manuscript.

## Competing interests

The authors declare no competing interests.

## Additional information

### Supplementary information

The online version contains supplementary material.

**Correspondence** and requests should be addressed to S.M.

## Notes

### Competing Interest Statement

The authors have declared no competing interest.

## References

1. Anderson GS, Barton PS, Archer M, Wallace JR. Invertebrate scavenging communities. In: Olea PP, Mateo-Tomás P, Sánchez-Zapata JA, editors. Carrion ecology and management. Cham: Springer International Publishing; 2019. p. 45–69.

2. Carter DO, Yellowlees D, Tibbett M. Cadaver decomposition in terrestrial ecosystems. Die Naturwissenschaften. 2007;94(1):12–24.

3. Barton PS, Cunningham SA, Lindenmayer DB, Manning AD. The role of carrion in maintaining biodiversity and ecological processes in terrestrial ecosystems. Oecologia. 2013;171(4):761–72.

4. Matuszewski S, Mądra-Bielewicz A. Competition of insect decomposers over large vertebrate carrion: Necrodes beetles (Silphidae) vs. blow flies (Calliphoridae). Curr Zool. 2021.

5. Dawson BM, Wallman JF, Evans MJ, Butterworth NJ, Barton PS. Priority effects and density promote coexistence between the facultative predator Chrysomya rufifacies and its competitor Calliphora stygia. Oecologia. 2022;199(1):181–91.

6. Verheggen F, Perrault KA, Megido RC, Dubois LM, Francis F, Haubruge E, et al. The odor of death: an overview of current knowledge on characterization and applications. Bioscience. 2017;67(7):600–13.

7. Dekeirsschieter J, Frederickx C, Lognay G, Brostaux Y, Verheggen FJ, Haubruge E. Electrophysiological and behavioral responses of Thanatophilus sinuatus Fabricius (Coleoptera: Silphidae) to selected cadaveric volatile organic compounds. J Forensic Sci. 2013;58(4):917–23.

8. Frederickx C, Dekeirsschieter J, Verheggen FJ, Haubruge E. Responses of Lucilia sericata Meigen (Diptera: Calliphoridae) to Cadaveric Volatile Organic Compounds. J Forensic Sci. 2012;57(2):386–90.

9. Nilssen AC, Tømmerås BÅ, Schmid R, Evensen SB. Dimethyl trisulphide is a strong attractant for some calliphorids and a muscid but not for the reindeer oestrids Hypoderma tarandi and Cephenemyia trompe. Entomol Exp Appl. 1996;79(2):211–8.

10. Kalinova B, Podskalska H, Ruzicka J, Hoskovec M. Irresistible bouquet of death--how are burying beetles (Coleoptera: Silphidae: Nicrophorus) attracted by carcasses. Die Naturwissenschaften. 2009;96(8):889–99.

11. Johansen H, Solum M, Knudsen GK, Hagvar EB, Norli HR, Aak A. Blow fly responses to semiochemicals produced by decaying carcasses. Med Vet Entomol. 2014;28(1):26–34.

12. Stensmyr MC, Urru I, Collu I, Celander M, Hansson BS, Angioy A-M. Rotting smell of dead-horse arum florets. Nature. 2002;420(6916):625–6.

13. Michaud JP, Schoenly KG, Moreau G. Rewriting ecological succession history: did carrion ecologists get there first? Q Rev Biol. 2015;90(1):45–66.

14. Dawson BM, Wallman JF, Evans MJ, Barton PS. Is resource change a useful predictor of carrion insect succession on pigs and humans? J Med Entomol. 2021;58(6):2228–35.

15. Charabidze D, Vincent B, Pasquerault T, Hedouin V. The biology and ecology of Necrodes littoralis, a species of forensic interest in Europe. Int J Legal Med. 2016;130(1):273–80.

16. Gruszka J, Krystkowiak-Kowalska M, Frątczak-Łagiewska K, Mądra-Bielewicz A, Charabidze D, Matuszewski S. Patterns and mechanisms for larval aggregation in carrion beetle Necrodes littoralis (Coleoptera: Silphidae). Animal Beh. 2020;162:1–10.

17. Matuszewski S, Konwerski S, Fratczak K, Szafalowicz M. Effect of body mass and clothing on decomposition of pig carcasses. Int J Legal Med. 2014;128(6):1039–48.

18. Löbl I, Löbl, Daniel,. Catalogue of Palaearctic Coleoptera. Volume 2: Hydrophiloidea –Staphylinoidea, Revised and updated edition. Leiden, Boston: Brill; 2015.

19. Ratcliffe BC. The natural history of Necrodes surinamensis (Fabr.)(Coleoptera: Silphidae). Trans AmEntomol Soc. 1972:359–410.

20. Ratcliffe BC. The carrion beetles (Coleoptera: Silphidae) of Nebraska 1996.

21. Charabidze D, Trumbo S, Grzywacz A, Costa JT, Benbow ME, Barton PS, et al. Convergence of social strategies in carrion breeding insects. BioScience. 2021.

22. Matuszewski S, Mądra-Bielewicz A. Heat production in a feeding matrix formed on carrion by communally breeding beetles. Front Zool. 2021;18(1):5.

23. Bajerlein D, Taberski D, Matuszewski S. Estimation of postmortem interval (PMI) based on empty puparia of Phormia regina (Meigen)(Diptera: Calliphoridae) and third larval stage of Necrodes littoralis (L.)(Coleoptera: Silphidae)–Advantages of using different PMI indicators. J Forensic Leg. Med. 2018;55:95–8.

24. Gruszka J, Matuszewski S. Temperature models of development for Necrodes littoralis L.(Coleoptera: Silphidae), a carrion beetle of forensic importance in the Palearctic region. Sci Rep. 2022;12(1):1–11.

25. Dekeirsschieter J, Frederickx C, Verheggen FJ, Boxho P, Haubruge E. Forensic entomology investigations from Doctor Marcel Leclercq (1924-2008): a review of cases from 1969 to 2005. J Med Entomol. 2013;50(5):935–54.

26. Lutz L, Zehner R, Verhoff MA, Bratzke H, Amendt J. It is all about the insects: a retrospective on 20 years of forensic entomology highlights the importance of insects in legal investigations. Int J Legal Med. 2021;135(6):2637–51.

27. Bonacci T, Mendicino F, Carlomagno F, Bonelli D, Scapoli C, Pezzi M. First report of the presence of Necrodes littoralis (L.)(Coleoptera: Silphidae) on a human corpse in Italy. J Forensic Sci. 2021;66(6):2511–4.

28. Dekeirsschieter J, Verheggen FJ, Gohy M, Hubrecht F, Bourguignon L, Lognay G, et al. Cadaveric volatile organic compounds released by decaying pig carcasses (Sus domesticus L.) in different biotopes. Forensic Sci Int. 2009;189(1-3):46–53.

29. Paczkowski S, Nicke S, Ziegenhagen H, Schütz S. Volatile emission of decomposing pig carcasses (Sus scrofa domesticus L.) as an indicator for the postmortem interval. J Forensic Sci. 2015;60:S130–S7.

30. Paczkowski S, Schütz S. Post-mortem volatiles of vertebrate tissue. Appl Microbiol Biotech. 2011;91(4):917–35.

31. Martin C, Verheggen F. Odour profile of human corpses: A review. Forensic Chemistry. 2018;10:27–36.

32. Statheropoulos M, Spiliopoulou C, Agapiou A. A study of volatile organic compounds evolved from the decaying human body. Forensic Sci Int. 2005;153(2-3):147–55.

33. Frederickx C, Dekeirsschieter J, Verheggen FJ, Haubruge E. Host-habitat location by the parasitoid, Nasonia vitripennis Walker (Hymenoptera: Pteromalidae). J Forensic Sci. 2014;59(1):242–9.

34. Trumbo ST, Steiger S. Finding a fresh carcass: bacterially derived volatiles and burying beetle search success. Chemoecol. 2020;30(6):287–96.

35. Matuszewski S, Bajerlein D, Konwerski S, Szpila K. Insect succession and carrion decomposition in selected forests of Central Europe. Part 3: Succession of carrion fauna. Forensic Sci Int. 2011;207(1-3):150–63.

36. Grassberger M, Frank C. Initial study of arthropod succession on pig carrion in a central European urban habitat. J Med Entomol. 2004;41(3):511–23.

37. Scott MP. The ecology and behavior of burying beetles. Annual Rev Entomol. 1998;43:595–618.

38. Matuszewski S. Estimating the pre-appearance interval from temperature in Necrodes littoralis L. (Coleoptera: Silphidae). Forensic Sci Int. 2011;212(1–3):180–8.

39. von Hoermann C, Ruther J, Reibe S, Madea B, Ayasse M. The importance of carcass volatiles as attractants for the hide beetle Dermestes maculatus (De Geer). Forensic Sci Int. 2011;212(1–3):173–9.

40. Matuszewski S, Bajerlein D, Konwerski S, Szpila K. Insect succession and carrion decomposition in selected forests of Central Europe. Part 2: Composition and residency patterns of carrion fauna. Forensic Sci Int. 2010;195(1-3):42–51.

41. Matuszewski S, Bajerlein D, Konwerski S, Szpila K. Insect succession and carrion decomposition in selected forests of Central Europe. Part 1: Pattern and rate of decomposition. Forensic Sci Int. 2010;194(1-3):85–93.

42. Whiteley M, Diggle SP, Greenberg EP. Progress in and promise of bacterial quorum sensing research. Nature. 2017;551(7680):313–20.

43. Ma Q, Fonseca A, Liu W, Fields AT, Pimsler ML, Spindola AF, et al. Proteus mirabilis interkingdom swarming signals attract blow flies. ISME J. 2012;6(7):1356–66.

44. Davis TS, Crippen TL, Hofstetter RW, Tomberlin JK. Microbial volatile emissions as insect semiochemicals. J Chem Ecol. 2013;39(7):840–59.

45. Tomberlin JK, Crippen TL, Wu G, Griffin AS, Wood TK, Kilner RM. Indole: an evolutionarily conserved influencer of behavior across kingdoms. Bioessays. 2017;39(2):1600203.

46. Cragg J, Thurston BA. The reactions of blowflies to organic sulphur compounds and other materials used in traps. Parasitology. 1950;40(1-2):187–94.

47. Brodie B, Gries R, Martins A, VanLaerhoven S, Gries G. Bimodal cue complex signifies suitable oviposition sites to gravid females of the common green bottle fly. Entomol Experiment Appl. 2014;153(2):114–27.

48. Zito P, Sajeva M, Raspi A, Dötterl S. Dimethyl disulfide and dimethyl trisulfide: so similar yet so different in evoking biological responses in saprophilous flies. Chemoecology. 2014;24(6):261–7.

49. Szpila K, Madra A, Jarmusz M, Matuszewski S. Flesh flies (Diptera: Sarcophagidae) colonising large carcasses in Central Europe. Parasitol Res. 2015;114(6):2341–8.

50. Bajerlein D, Matuszewski S, Konwerski S. Insect succession on carrion: Seasonality, habitat preference and residency of histerid beetles (Coleoptera: Histeridae) visiting pig carrion exposed in various forests (Western Poland). Polish J Ecol. 2011;59(4):789–97.

51. Anton E, Niederegger S, Beutel RG. Beetles and flies collected on pig carrion in an experimental setting in Thuringia and their forensic implications. Med Vet Ent. 2011;25(4):353–64.

52. Frederickx C, Dekeirsschieter J, Brostaux Y, Wathelet JP, Verheggen FJ, Haubruge E. Volatile organic compounds released by blowfly larvae and pupae: New perspectives in forensic entomology. Forensic Sci Int. 2012;219(1–3):215–20.

53. Faul F, Erdfelder E, Lang A-G, Buchner A. G* Power 3: A flexible statistical power analysis program for the social, behavioral, and biomedical sciences. Behav Res Meth. 2007;39(2):175–91.

